# High throughput hemogram of T cells using digital holographic microscopy and deep learning

**DOI:** 10.1101/2021.12.23.473983

**Authors:** Roopam K. Gupta, Nils Hempler, Graeme P. A. Malcolm, Kishan Dholakia, Simon J. Powis

**Affiliations:** Atonarp Micro-Systems India Pvt. Ltd., The Millenia, Tower A, 3rd Floor, No. 1 & 2 Murphy Road, Ulsoor, Bangalore 560008 India; School of Medicine and Biomedical Sciences Research Complex, University of St. Andrews, KY16 9TF, UK; SUPA, School of Physics and Astronomy, University of St. Andrews, KY16 9SS, UK; Department of Physics, College of Science, Yonsei University, Seoul 03722, South Korea; M Squared Lasers, 1 Kelvin Campus, West of Scotland Science Park, Glasgow, G20 OSP, UK

**Keywords:** Deep learning, Microscopy, Immunology, Multivariate analysis

## Abstract

T cells of the adaptive immune system provide effective protection to the human body against numerous pathogenic challenges. Current labelling methods of detecting these cells, such as flow cytometry or magnetic bead labelling, are time consuming and expensive. To overcome these limitations, the label-free method of digital holographic microscopy (DHM) combined with deep learning has recently been introduced which is both time and cost effective. In this study, we demonstrate the application of digital holographic microscopy with deep learning to classify the key CD4^+^ and CD8^+^ T cell subsets. We show that combining DHM of varying fields of view, with deep learning, can potentially achieve a classification throughput rate of 78,000 cells per second with an accuracy of 76.2% for these morphologically similar cells. This throughput rate is 100 times faster than the previous studies and proves to be an effective replacement for labelling methods.

## 1. Introduction

The adaptive immune response comprises white blood cells including T and B cells that can recognise and respond in an antigen-specific manner to a vast array of potential human pathogens. Of great significance, residing within this same subset of cells is the ability to generate memory cells, which can produce faster and stronger secondary responses. Vaccination/immunisation relies almost exclusively on the generation of such memory T and B cells to protect both individuals and populations [1].

T cells at the most basic level of functionality are divided into two groups based upon their expression of CD4 and CD8 cell surface proteins [2]. Typically, CD4^+^ T cells coordinate both B cell antibody responses and other T cells by the secretion of various cytokines [3], coordinated through expression of HLA class II, whereas CD8^+^ T cells are usually directly capable of elimination of virally infected or tumourigenic cells by the detection of specific viral or tumour antigenic peptides presented on HLA class I molecules [4]. The numbers of T cells can vary significantly during the course of diseases. For example, in HIV the numbers of CD4^+^ T cells can reduce to very low levels over time [5], and recent data for patients with COVID-19 has shown loss of both CD4^+^ and CD8^+^ population in many patients undergoing ICU-level care [6, 7, 8].

The identification of these cells requires destructive fixation or chemical staining which is both time consuming and costly. To circumvent these issues, label-free optical methods of Raman spectroscopy [9], autofluorescence lifetime imaging [10] or digital holographic microscopy (DHM) have been employed [11]. These methods provide molecular or morphological data and require an additional step of statistical analysis. Methods such as support vector machines (SVMs), random forests (RFs) or artificial neural networks (ANNs) have been popularly employed for these purposes. However, due to the inherent linearity, the methods of SVM and RF have proven to be less efficient for classification than deep learning based ANNs [12]. Hence deep learning is being ever more widely applied to solve the classification problem in biophotonics [13]. Another aspect of the mentioned systems is their throughput rate. While Raman spectroscopy provides high molecular specificity, it is slow and lacks the aspect of throughput [14]. DHM when combined with convolutional neural networks, on the other hand, provides the capability to differentiate morphologically similar cells with a recent demonstration of throughput rate of more than 100 cells/s [15]. This throughput rate is still too low: the gold standard flow cytomtery may allow a throughput rate of 70,000 cells/s [16].The throughput rate of the DHM system can be enhanced by reducing the magnification and numerical aperture (NA) of the microscopic objective which may in-turn result in a lower resolution of images. These lower resolution images system can be transformed into ones of higher resolution using the single image super resolution (SISR) method of deep learning [17]. Recently, deep learning has been widely applied more broadly in photonics to improve the resolution of bright field optical microscope [18], to enable cross-modality super resolution in fluorescence microscopy [19], to facilitate pixel super-resolution in coherent imaging systems Liu et al. [20], and to enhance the resolution of scanning electron microscopy [21].

Here, we address the use of DHM for rapid, high throughput classification of CD4^+^ and CD8^+^ T cells. We present a method based upon particle swarm optimization (PSO) [22] to identify an optimal CNN geometry for a given dataset. Subsequently, we compare the classification performance of DHM-CNN combination for different optical magnifications. We also present a new method of SISR in microscopy based on cycle generative adversarial networks (GANs) for the enhancement in the resolution of images acquired from 20X optical magnification to images acquired from 100X optical magnification. Compared to previous studies [19, 20, 21], which require an additional step of co-registering the field of view (FOV), our semi-supervised method improves the resolution of unpaired phase images which were independent of FOV and do not require any additional analytical methods. Our approach demonstrates a possibility of high throughput of 78,000 cells/s using a combination of DHM with CNNs, which is nearly two orders of magnitude in excess of previous reports [15]. Importantly, this result for the first time makes a label-free DHM approach comparable to the gold standard of flow cytometry.

## 2. Methods

### 2.1. Cell Isolation

This work was undertaken after ethical review from the School of Medicine at the University of St. Andrews, utilising buffy coats of six different healthy donors obtained from NHS UK. PBMC were isolated from the buffy coats by centrifugation at room temperature on Ficoll-Pacque at density 1.077 g/ml (Thermofisher, UK). CD4^+^ and CD8^+^ T cell populations were isolated using a negative depletion method following the manufacturers instructions (Dynabeads CD4^+^ T cells, 11346D and Dynabeads CD8^+^ T cells, 11348D, Thermofisher UK). Following the isolation, the purified cells were cultured in RPMI 1640 supplemented with 5 % Foetal Bovine Serum (both Thermofisher, UK).

Flow cytometry was employed to confirm the purity of the purified cell samples. Each cell type was stained with combinations of antibodies CD3-PE and -FITC, clone HIT3a, eBioscience UK, CD4-PE and -FITC, clone OKT4, eBioscience UK, CD8-PE and -AF488, clones SK1, eBioscience UK, and FAB1509G, R&D UK. Cells were analysed on a Guava 8HT cytometer (Merck Millipore, UK).

For the optical analysis, the cells were resuspended in Phosphate Buffer Saline (PBS) with 0.5% FBS solution to avoid aggregation. 20 *μ*l of cell suspension was transferred to the center of a clean quartz slide (25.4 mm × 25.4 mm × 1 mm) chamber - formed by a 100 *μ*m thick vinyl spacer. This chamber was covered from the top using a thin quartz slide (25.4 mm × 25.4 mm, 0.11 mm - 0.15 mm thickness) and finally the whole assembly was inverted and left for ∼20 minutes to avoid cellular motion.

### 2.2. Digital holographic microscopy

We modified a previously employed off-axis digital holographic microscope to capture the holographic images of the cells with three different optical configurations [15]. As shown in Fig. 1, the optical configurations were varied by changing the microscopic objectives to ones with magnifications of 20X (NA = 0.4, Nikon Japan 130314), 60X (NA = 0.8, Nikon) and 100X (NA = 0.9, Nikon Japan 230538) respectively. The sample was placed between the two objectives and the image was interfered with the light from the reference arm at the surface of the CCD camera (Ximea XiQ MQ013MG-E2). This camera was set to accumulate 16 bit images with a frame rate of 60 fps and an exposure time of 33.3 ms.

**Figure 1:**
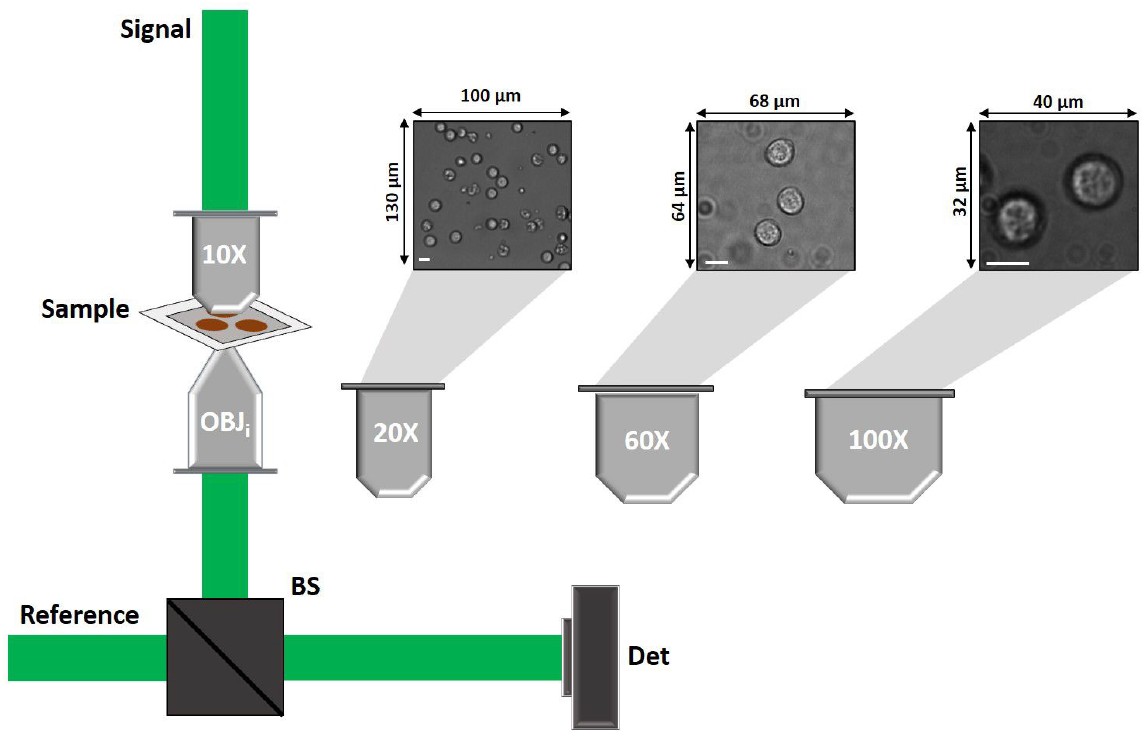
Schematic of different optical configurations used with digital holographic microscope. The three objectives with the magnifications of 20X, 60X and 100X were employed for data acquisition. The sub figures indicate three bright field FOVs acquired for each objective. The scale bar indicates 10 *μ*m for each image.

We acquired the data for the two cell types separately for each optical configuration. The data acquired using the three objectives varied with the magnification. With an increase in magnification, the field of view (FOV) was reduced. For the 20X objective (lateral resolution: 0.66 *μ*m; axial resolution: 6.65 *μ*m), we achieved an FOV of 100 *μ*m × 130 *μ*m; for the 60X objective (lateral resolution: 0.31 *μ*m; axial resolution: 1.47 *μ*m), we achieved 68 *μ*m × 64 *μ*m for the FOV and 100X objective (lateral resolution: 0.29 *μ*m; axial resolution: 1.31 *μ*m) allowed for an FOV of 40 *μ*m × 32 *μ*m. Since these FOVs provide with cell images of different radii (in px), we implemented Haugh transform based method.

### 2.3. k-means segmentation

The phase images calculated using the three optical configurations demonstrate different degrees of resolutions. Hence to identify and understand the degree of granularity discovered using each configuration, we implemented a method of k-means clustering based image segmentation [23]. We considered the phase images corresponding to each configuration individually. To identify the number of segmentation classes across the cellular structure, we increased the number of segmentation classes in steps of unity until the algorithm returned a solution with discontinuous boundaries.

In this specific case of DHM, as the phase images represent the refractive index variation across the image, the classes represent this distribution. This in turn demonstrates the variation of granularity analyzed using the phase images evaluated using each FOV.

### 2.4. Optimization of the CNN geometry

The phase images obtained using the three configurations were of different sizes, for the 20X objective the phase images were 52 × 52 pixels (px) whereas for the 60X objective, the phase images were of size 100 × 100 px and for 100X objective the phase images were acquired with a size of 200× 200 px. We optimized a CNN geometry for each of the three configuration by implementing a PSO based approach. PSO is a type of swarm intelligence method for global optimization where each individual (called particle) of the population (called swarm) adjust their trajectory towards the previous best position attained by any member of their topological neighborhood. This approach is used to minimize the error output of an objective function. In our case, we consider the objective function as the classification sensitivity and specificity achieved using a given network geometry (particle).

To identify the best CNN geometries, we divided the complete dataset for each optical configuration into training, validation and test sets such that the training/validation set and testing set came from different donors. Details of segmentation of above datasets have been summarised in table 1.

**Table 1:** Table summarizing the total number of single cells phase images considered for different optical configurations.

| S.No. | 20X |  | 60X |  | 100X |  |
| --- | --- | --- | --- | --- | --- | --- |
|  | CD4 | CD8 | CD4 | CD8 | CD4 | CD8 |
| **Train/Val** | 2385 | 2056 | 1066 | 971 | 704 | 704 |
| **Test** | 344 | 323 | 84 | 77 | 104 | 96 |

We implemented the PSO algorithm by constructing an objective function in the form of a training instance. Each training instance was designed to develop a network geometry and providing the performance of the geometry on the validation dataset. For each training instance, we trained the CNNs using an Adam optimizer [24] with maximum epochs set at 100, initial learning at 1 × 10^−3^, L2 regularization at 5 × 10^−6^, validation frequency at 40 iterations and validation patience of 5 iterations with a mini batch size of 128 images. The network geometry was developed by the virtue of parameters in the form of each particle in the PSO algorithm. These parameters dictated the number of layers, type of layers, number of filters in each layer, stride and padding for each layer. To conserve maximal input image information, the convolution layers were restricted with filter sizes between 1 and 5. To conserve the memory of system and avoid over estimation, number of convolution filters for any layer were restricted to a maximum of 50. The number of neurons in fully connected layer and the dropout ratio were restricted to be more than zero. To conserve the network geometry, the input layer was set as image input layer with the size of image in the dataset and the output layer was set with fully connected layer with 2 neurons (representing each class) followed by softmax layer and a classification layer. We evaluated the cost function for each training instance as:

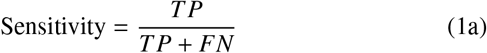

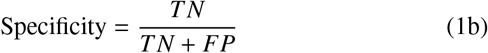

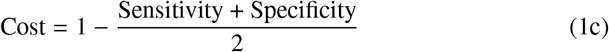

Here, in Eq. 1a and 1b, TP is true positive, TN is true negative, FP is false positive and FN is false negative.

We considered a total of 40 particles and a single swarm (optimized from 2 to 60 in the steps of one unit to avoid divergence) to find isolate an optimal architecture of CNN for each image size. Each particle’s position and velocity were initialized randomly. After the calculation of cost for all the particles, the particle with least cost was considered as the reference such that the position and velocity of all the other particles were updated relatively to the reference.

### 2.5. Cycle generative adversarial training for image transformation

The phase images evaluated using the three optical configuration show variability in the resolution due to different resolving powers of the microscopic objectives. As mentioned in section 2.2, the phase images evaluated from the 20X objective show the least resolution whereas the images captured using 100X objective display highest resolution. Hence to gather high resolution images with high throughput rate, we considered training a CNN to transform the images acquired using 20X objective into images acquired using 100X objective. In the current DHM system, it is very challenging to identify same cells using two different configurations, hence we trained the deep networks on unpaired images using cycle-generative adversarial training [25, 26].

As shown in Fig. 2, we developed two CNNs such that the input image could be down-sampled and then up-sampled to a required size at the output. For the transformation of phase images captured using 20X optical configuration to 100X optical configuration, we developed and optimized a 54 layered CNN (𝒢_20*X*→100*X*_: *=*𝒢_*a*_) whereas for the inverse translation, we developed a 34 layered CNN (𝒢 _100*X*→20*X*_ :*=*𝒢_*b*_). These networks were optimized by changing the network filters in the step of 8 units with respect to their performance on validation dataset. With respect to the training module requirement, we also developed and two discriminator networks with 23 layers (𝒟_100*X*_*:=*𝒟_*a*_) and 15 layers (𝒟_20*X*_*:=*𝒟_*b*_) respectively. We trained these networks with 300 randomly selected phase images each of CD4^+^ and CD8^+^ T cells from both the 20X and 100X optical configurations. Out of these, we considered 225 images for training and 75 images for validation.

**Figure 2:**
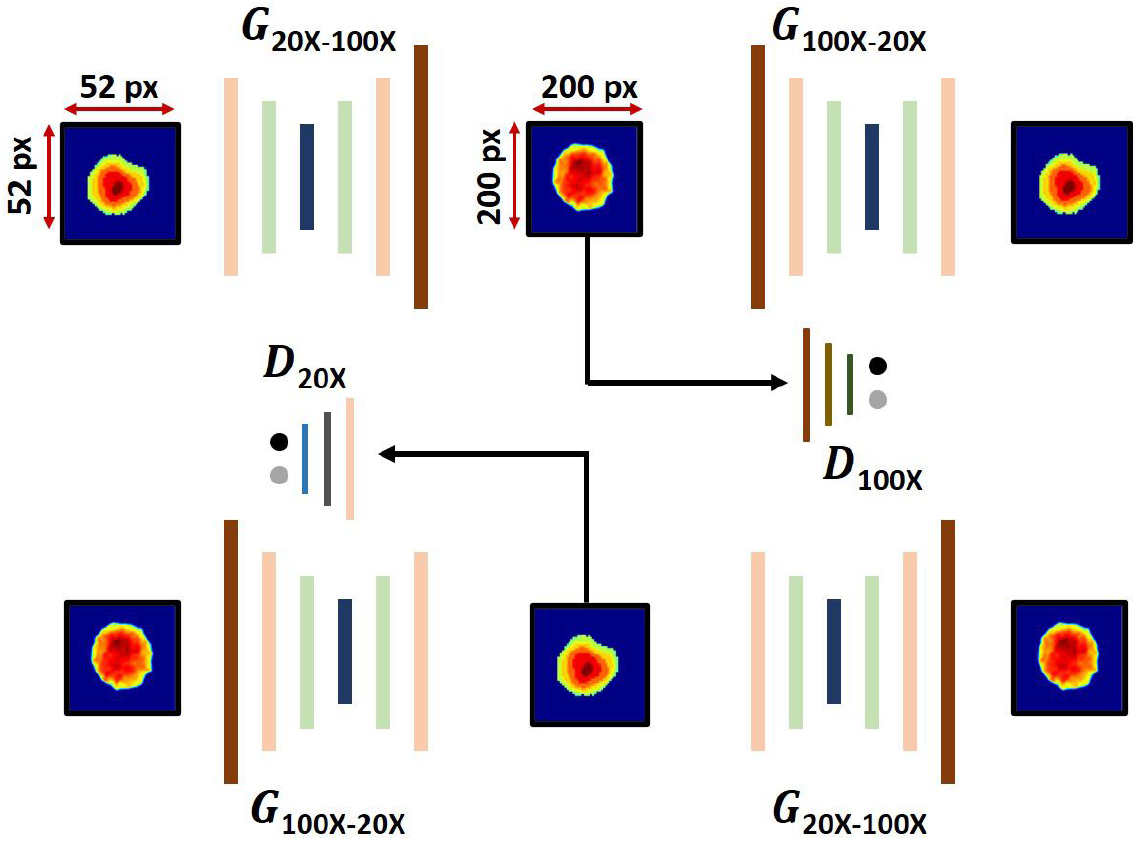
Schematic of cycle GAN model applied for super-resolving the phase images. The generative models 𝒢_20*X*→100*X*_ and 𝒢_100*X*→20*X*_ are trained with in a cycle consistent manner such that the inverse transformation of the images is conserved.

For each training instance, we calculated a generative adversarial loss for both the generative networks. We also calculated a cycle consistency loss using the combination of two networks. The generative adversarial loss was evaluated as:

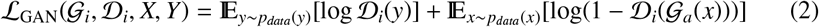

Here, *i* ∈ {*a, b*}. The cycle consistency loss ℒ_cyc_(𝒢_*a*_, 𝒢_*b*_) is computed, to satisfy the condition *x* → 𝒢_*a*_(*x*) → 𝒢_*b*_(𝒢_*a*_(*x*)) ≈ *x*, as:

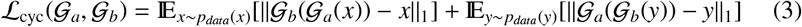

Here, the variables *x* and *y* represent the input and output images for the given network configuration. The combined loss was calculated as:

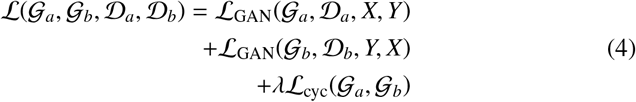

Here, *λ* is a hyperparameter which we chose as 10 for this application. During the training, the objective is to minimize the combined loss for the generator networks while maximizing the loss for the discriminator networks:

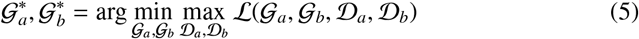

To achieve minimum training loss and avoid divergence during training, we considered the training batch images in the mini-batches of 45 images. An Adam optimizer was considered with a learning rate of 2 × 10^−4^, gradient descent factor of 0.5 and a squared gradient descent factor of 0.99. We validated the network performance after 25 iterations using 25 randomly sampled images for both the cell types from the validation set. The training was continued for a total of 5000 epochs.

## 3. Results

### 3.1. Cell isolation

Untouched human blood CD4^+^ and CD8^+^ T cells were obtained by negative depletion, in which other cells not of interest were removed using cell-lineage specific antibodies. Flow cytometry (Fig. 3) confirmed the purity of the cell populations in line with our previous studies [27, 28] with CD4^+^ cells isolated at an average of 89% (n=3) and CD8^+^ cells at an average of 86% (n=3).

**Figure 3:**
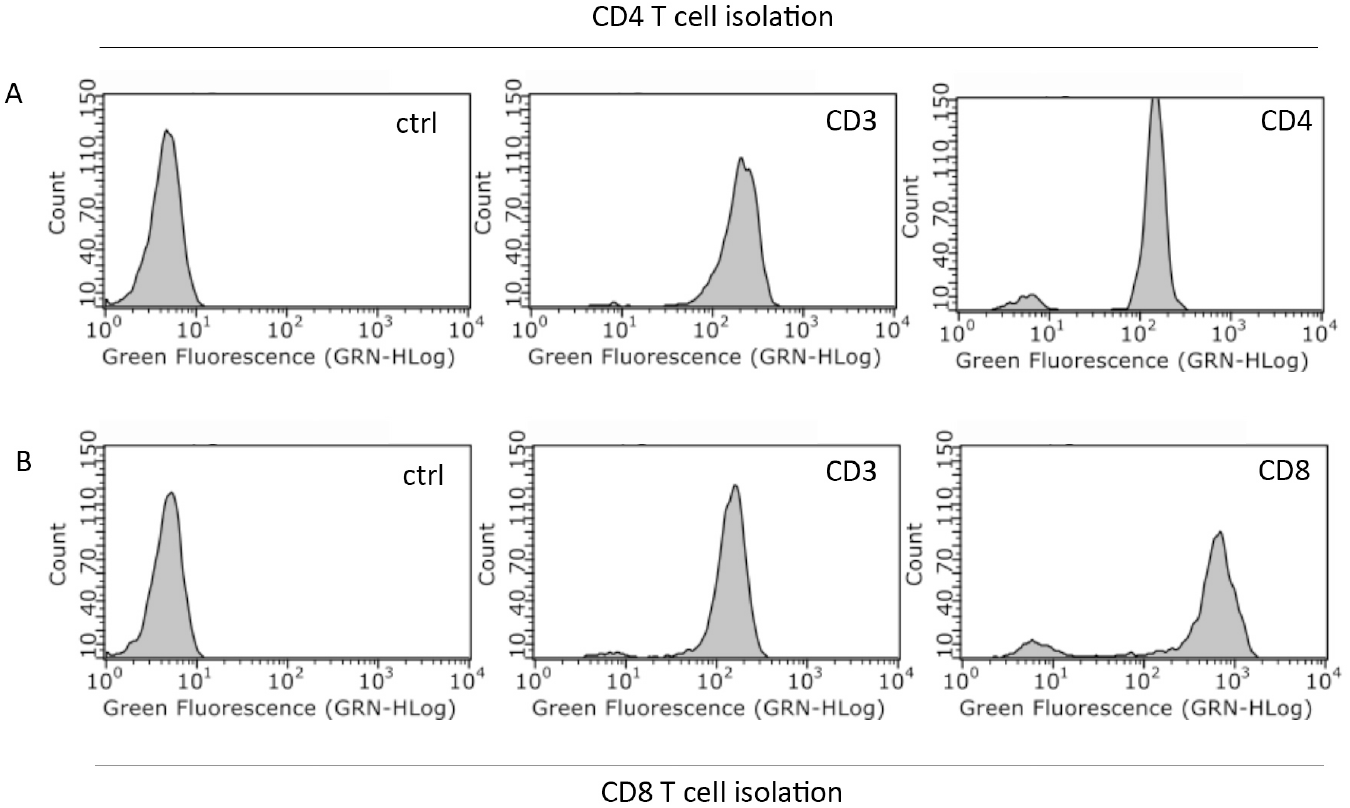
Representative flow cytometric plots of CD4 and CD8 T cells purified by negative depletion. Purified cell samples were stained with anti-CD3, -CD4 or –CD8-FITC or AF488 coupled antibodies and analysed by flow cytometry for (A) CD4 T cells and (B) CD8 T cells. Average purity of three separate purifications is reported in the main text.

### 3.2. Automated detection of cells and phase image calculation

We captured bright field and fringe images using all the three configurations described above. These images presented with variable radii of single cells, hence to automatically detect these cells, we implemented Haugh transform using a prewritten MATLAB script [29].

We optimized the search parameters of radii, gradient threshold and radius of search filter with respect to the images acquired for each configuration. In order to optimize these parameters, we considered the size of cells, mean magnitude of gradient for empty space and the radius of cells for each configuration.

The images accumulated using the three optical configuration exhibit varying FOVs and resolutions. Fig. 4 demonstrate the automatic cellular detection for various FOVs. As summarised in Table 2, the FOV achieved by using the 20X objective was greatest at 100 *μ*m × 130 *μ*m (which may allow imaging a maximum of 1300 cells in one snapshot which potentially allowed a throughput rate of 78,000 cells per second), however, the resolution of the accumulated images was poor. For imaging using the 60X objective, a smaller FOV of 68 *μ*m × 64 *μ*m was achieved (allowing a maximum of 36 cells resulting in the highest possible throughput of 2,160 cells per second) with a reasonable resolution. Imaging using the 100X objective resulted in a much smaller FOV of 40 *μ*m × 32 *μ*m (enclosing a maximum of 12 cells and allowing a highest possible throughput of 720 cells per second) with the highest resolution.

**Table 2:** Summary of field of views and maximum realizable throughput for the three optical configurations.

| Optical Configuration | Field of View | Realizable Throughput |
| --- | --- | --- |
| 20X | $100 \times 130 \mu\text{m}$ | 78000 cells/s |
| 60X | $68 \times 64 \mu\text{m}$ | 2160 cells/s |
| 100X | $40 \times 32 \mu\text{m}$ | 720 cells/s |

**Figure 4:**
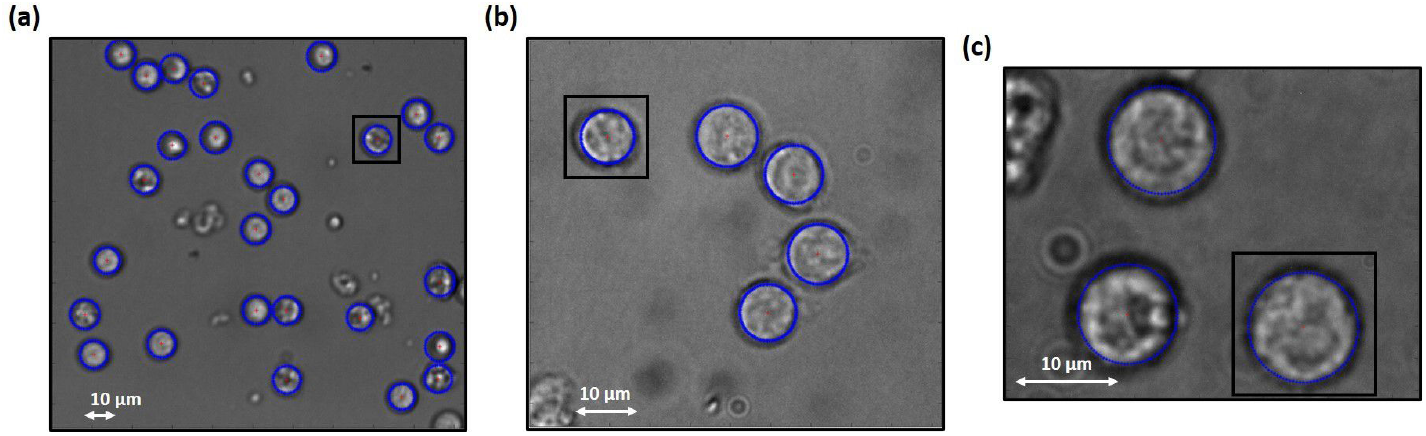
Automatic detection of cells using Haugh transform. Subsection of Bright field images recovered from (a) 20X Objective (100 *μ*m × 130 *μ*m) (b) 60X objective (68 *μ*m × 64 *μ*m) and (c) 100X objective (40 *μ*m × 32 *μ*m). Blue highlighted regions represent the automatic detection of cells for three FOV’s using Haugh transform circular detection. Here the boxes show the cropped area of images for single cells.

With respect to the numerical aperture of the microscopic objectives, the retrieved phase images show the differences in resolution. The phase image recovered using 20X objective with a numerical aperture (NA) of 0.4, displays the least resolution (lateral: 0.66 *μ*m; axial: 6.65 *μ*m) for both the cell lines (Fig. 5 (a),(d)). The application of 60X objective with the NA of 0.8 results in moderately resolved (lateral resolution: 0.31 *μ*m; axial resolution: 1.47 *μ*m) phase images (Fig. 5 (b),(e)) and the phase images calculated from the fringe images captured using the 100X objective (with the NA of 0.9) were highly resolved (lateral resolution: 0.29 *μ*m; axial resolution: 1.31 *μ*m).

**Figure 5:**
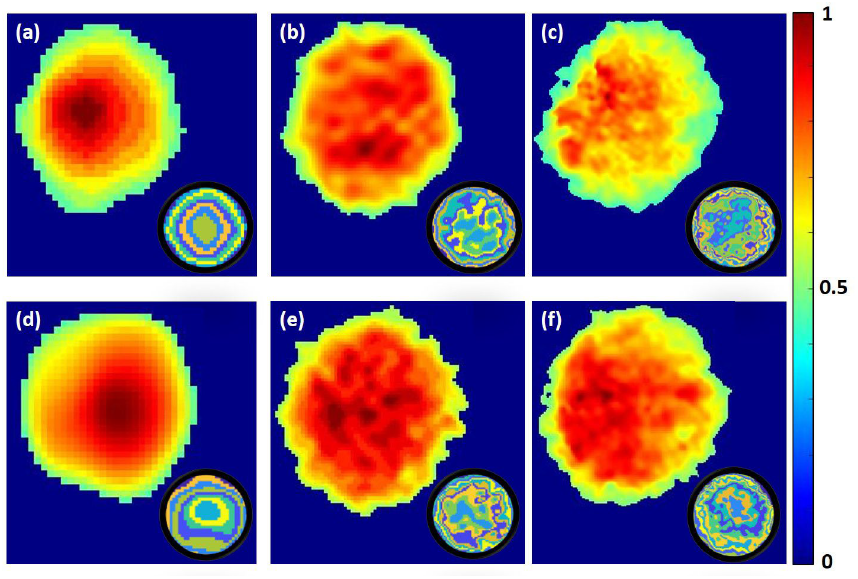
Normalized phase images of T Cells. Single cell normalized phase images of the CD4 cells retrieved using (a) 20X objective (b) 60X objective (c) 100 X objective; Single cell normalized phase images of CD8 cells retrieved using (d) 20X objective (e) 60X objective (f) 100X objective. Colorbar represents the normalized phase gain of the signal arm with respect to the reference arm. Here the sub figures are the k-means based image segmentation of these images.

After the extraction of single cell phase images, we employed k-means clustering based segmentation to quantify the granularity achieved using different configurations. As anticipated, the phase images of the two cell types show a very similar variation in resolution with respect to the objectives. As shown in sub-figures of Fig. 5, the algorithm when applied over the phase images of the two cells for the 20X objective, saturated at 8 segments. For the phase images accumulated using 60X objective, the algorithm saturated at 9 segments for CD4 cells whereas it saturated at 10 segments for the CD8 cells. For its application on phase images evaluated using 100X objective, the algorithm presented a saturation at 11 segments for both the cell types.

These results clearly demonstrate that the images acquired using the three objectives show a variability in resolution. It is also evident that with an increase in the numerical aperture of the objective, these images provide the information of a wider range of variations across the cellular structure.

### 3.3. Classification of phase images

The next step for the analysis of the single phase images was to classify them with respect to the cell types. This was achieved by employing the CNNs which were optimized by implementing PSO algorithm as explained before.

The three optical configurations, resulted in different sizes of single cell phase images as 52 × 52 px for 20 X objective, 100 × 100 px for 60 X objective and 200 ×200 px for 100X objective. Hence the optimal CNN geometry also displayed a variation in size. For the 20X optical configuration, the optimal CNN geometry was identified with a total of six layers with 39,998 parameters. Using the validation set, the CNN returned a sensitivity of 63.13 % ± 2.23 % and specificity of 64.93% ±5.65%, whereas when considered for the test dataset, the CNN resulted in a sensitivity of 64.07 % ± 2.64 % and a specificity of 56.83 % ±2.36 %.

For the 60X optical configuration, the optimal CNN geometry was identified as a slightly longer network. This geometry comprised a total of 16 layers with 226,707 parameters. On the validation set, the classification efficiency of the CNN resulted in 70.94 % ±2.27 % sensitivity and 65.52 % ±2.08% specificity, whereas on the test set the trained CNN resulted in a specificity of 69.92 % ± 3.91% and a sensitivity of 69.59 % ±3.10 %. Finally, the optimization routine was also implemented on the phase images acquired using the 100X optical configuration and this resulted in a CNN geometry of 24 layers with 1,603,327 parameters. This geometry when applied over the phase images from validation dataset resulted in the specificity of 80.28 % ±1.17 % and a sensitivity of 77.77 % ± 2.72 %. The specificity and sensitivity calculated using the test data were 82.5 % ±3.96 % and 73.18 % ± 7.55 % respectively.

The interesting aspect of this comparison is the trend of increasing classification accuracy (Fig. 6) and decreasing throughput rate with respect to the optical configuration. For the 20X configuration, the optimal CNN geometry resulted in a validation accuracy of 62.28 % ± 2.54 % and 59.43 % ± 1.99 % as the test accuracy with an allowed maximum throughput rate of 78,000 cells per second. For the optical configuration with 60X magnification microscopic objective, the optimum CNN resulted in 67.98 % ± 0.27 % validation accuracy and 69.31 % ±2.04 % test accuracy. While considering the dataset accumulated using the microscopic objective with 100X magnification, the optimal CNN geometry resulted in a maximum classification accuracy with 78.91 % ±1.57 % for validation set and 76.2 % ± 5.27 % for the test set. These results confirm that by increasing the magnification of a holographic system, one may acquire an increasingly more precise classification of immune cells. However, with increasing magnification the throughput limit of the system deteriorates. Hence to keep an increased throughput limit and simultaneously improving the classification ability of the system, we trained another deep learning based model to transform the phase images acquired from 20X configuration to the phase images which may represent the acquisition from 100X optical configuration.

**Figure 6:**
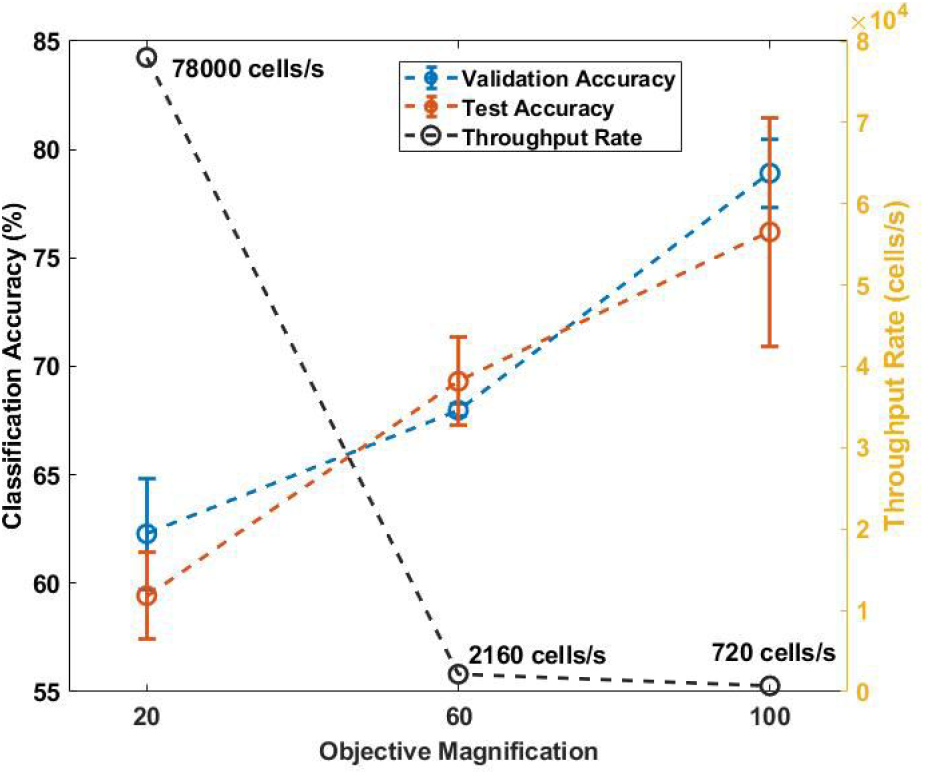
Comparison between throughput rate and classification accuracy obtained using different optical configurations. For the validation set, the values for classification accuracy range from 62.28 % ± 2.54 % for 20X objective, 69.31 % ± 2.04 % for 60X objective and 78.91 % ± 1.57 % for 100X objective. For the test set, a classification accuracy of 59.43 % ± 1.99 % for 20X objective, 69.31 % ± 2.04 % for 60X objective and 76.2 % ± 5.27 % for 100X objective was obtained. The three optical configurations allow for 78000 cells/s, 2160 cells/s and 720 cells/s as the throughput rate for 20X, 60X and 100X objective respectively. Here the curve in blue represents classification accuracy for validation set, curve in red represents the accuracy for test set and curve in black shows the variation in throughput rate for the three configurations.

### 3.4. Image transformation and classification

The single image super resolution transformation was implemented on the phase images acquired using 20X configuration to convert them into the phase images acquired using 100X configuration. To achieve this the DL models were trained using the cycle GAN training method as explained before.

The trained generative models resulted in astounding transformations of the phase images. As shown in Fig. 7, the trained deep generative model transformed the phase images with a high speed of 6.89 milliseconds per transformation (limited by the the processing power of the computer which can further be improved). In the mentioned figure, (a) shows the transformation of CD4^+^ T cells from 20X configuration into 100X configuration; (b) demonstrates the same transformation of CD8^+^ T cells. An interesting aspect which is visible on the transformed images is that the deep models automatically learned to draw an outline around the periphery of the cells. Another aspect which is evidently visible from these transformations is the variation in the shape of the cells. This can be explained with respect to the cycle GAN type training module. The CNNs trained with this training module learn to transform and simultaneously inverse transform the images between the two domains. This learning process makes sure that the statistics and the functional relationship between the two domains are maintained.

**Figure 7:**
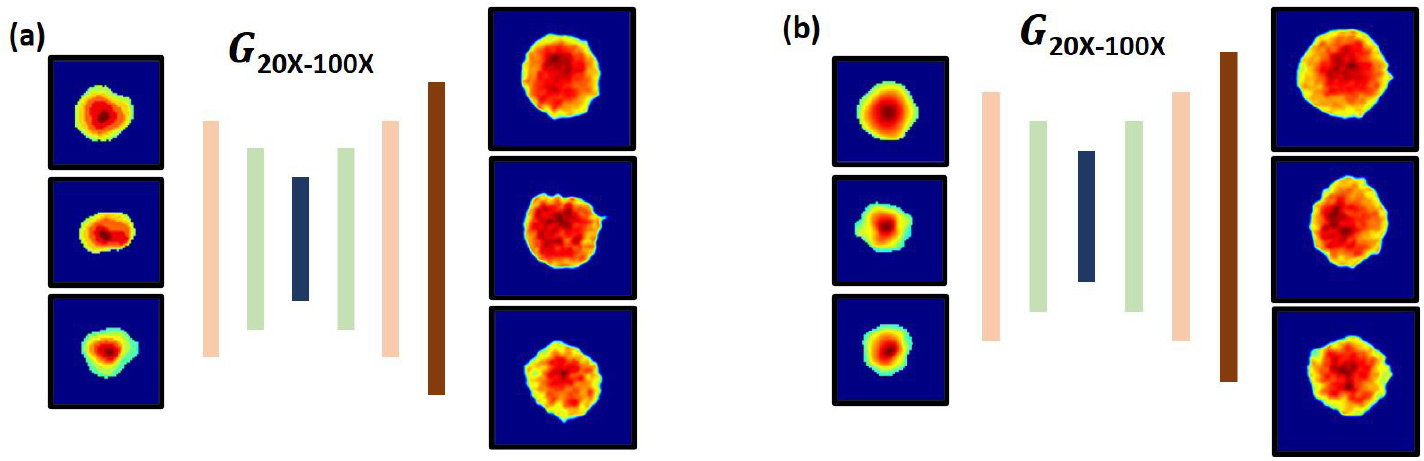
Demonstration of image transformation using the trained deep generative model. Transformation of phase images of (a) CD4^+^ and (b) CD8^+^ T cells acquired from 20X optical configuration into 100X optical configuration.

After the transformation of the phase images, we performed a classification using the pre-trained optimal CNN geometry. This resulted in the classification accuracy of 45.93 % ± 2.46 % for the validation set and 47.91 % ±2.05 % for the test set. These values were below the expectation and which may be explained due to the presence of boundaries and overall shape orientation of the cells. Hence, to overcome these problems, we re-trained the previously trained networks on the transformed image dataset. This resulted in satisfactory results with sensitivity of 81.55 % ±0.81 % and a specificity of 84.72 % ±1.16 % for the validation set and, the sensitivity and specificity of 79.77 % ± 3.32 % and 81.77 % ± 1.51 % for test set respectively.

These results display the increased capability in terms of classification accuracy for the 20X optical configuration. Hence, the application of cycle generative models would be beneficial for improving the resolution of system and simultaneously improving the throughput rate by up to two orders of magnitude.

## 4. Conclusion

In conclusion, we have presented a comparative study for the label free classification of T-cell subsets namely CD4^+^ and CD8^+^ T cells using a combination of digital holographic microscopy and convolutional neural networks. We compare the performance of DHM - CNN based classification by considering three different optical configurations. These configurations were considered by changing the optical magnification of the microscopic objectives between 20X, 60X and 100X. The T - cell subsets, being morphologically very similar, makes it very challenging for classification using the CNNs. Hence we report a maximal classification accuracy of 76.2% by using a microscopic objective with 100X magnification. Additionally, we demonstrate that the application of cycle GAN type training may help in enhancing the throughput rate and resolution of a DHM based system by up to two orders of magnitude.

## Author Contributions

KD and SJP developed the project. RKG performed the experiments and developed the numerical analysis procedures. RKG wrote the paper with contributions from SJP and KD which was approved by NH and GPAM. KD, SJP and NH supervised the project.

## References

[1] B. Pulendran, R. Ahmed, Immunological mechanisms of vaccination, Nature immunology 12 (2011) 509.

[2] W. Ellmeier, S. Sawada, D. R. Littman, The regulation of cd4 and cd8 coreceptor gene expression during t cell development, Annual review of immunology 17 (1999) 523–554.

[3] K. A. Read, M. D. Powell, B. K. Sreekumar, K. J. Oestreich, In vitro differentiation of effector cd4+ t helper cell subsets, in: Mouse Models of Innate Immunity, Springer, 2019, pp. 75–84.

[4] A. M. Van der Leun, D. S. Thommen, T. N. Schumacher, Cd8+ t cell states in human cancer: insights from single-cell analysis, Nature Reviews Cancer (2020) 1–15.

[5] G. Doitsh, W. C. Greene, Dissecting how cd4 t cells are lost during hiv infection, Cell host & microbe 19 (2016) 280–291.

[6] N. P. Restifo, M. E. Dudley, S. A. Rosenberg, Adoptive immunotherapy for cancer: harnessing the t cell response, Nature Reviews Immunology 12 (2012) 269–281.

[7] D. M. Pardoll, The blockade of immune checkpoints in cancer immunotherapy, Nature Reviews Cancer 12 (2012) 252–264.

[8] B. Diao, C. Wang, Y. Tan, X. Chen, Y. Liu, L. Ning, L. Chen, M. Li, Y. Liu, G. Wang, et al., Reduction and functional exhaustion of t cells in patients with coronavirus disease 2019 (covid-19), Frontiers in Immunology 11 (2020) 827.

[9] M. Chen, N. McReynolds, E. C. Campbell, M. Mazilu, J. Barbosa, K. Dholakia, S. J. Powis, The use of wavelength modulated raman spectroscopy in labelfree identification of t lymphocyte subsets, natural killer cells and dendritic cells, PLoS One 10 (2015) e0125158.

[10] A. J. Walsh, K. P. Mueller, K. Tweed, I. Jones, C. M. Walsh, N. J. Piscopo, N. M. Niemi, D. J. Pagliarini, K. Saha, M. C. Skala, Classification of t-cell activation via autofluorescence lifetime imaging, Nature Biomedical Engineering (2020) 1–12.

[11] N. McReynolds, F. G. Cooke, M. Chen, S. J. Powis, K. Dholakia, Multimodal discrimination of immune cells using a combination of raman spectroscopy and digital holographic microscopy, Scientific reports 7 (2017) 43631.

[12] E. Raczko, B. Zagajewski, Comparison of support vector machine, random forest and neural network classifiers for tree species classification on airborne hyperspectral apex images, European Journal of Remote Sensing 50 (2017) 144–154.

[13] P. Pradhan, S. Guo, O. Ryabchykov, J. Popp, T. W. Bocklitz, Deep learning a boon for biophotonics?, Journal of Biophotonics 13 (2020) e201960186.

[14] L. Woolford, M. Chen, K. Dholakia, C. S. Herrington, Towards automated cancer screening: label-free classification of fixed cell samples using wavelength modulated raman spectroscopy, Journal of biophotonics 11 (2018) e201700244.

[15] R. K. Gupta, M. Chen, G. P. Malcolm, N. Hempler, K. Dholakia, S. J. Powis, Label-free optical hemogram of granulocytes enhanced by artificial neural networks, Optics express 27 (2019) 13706–13720.

[16] J. Picot, C. L. Guerin, C. Le Van Kim, C. M. Boulanger, Flow cytometry: retrospective, fundamentals and recent instrumentation, Cytotechnology 64 (2012) 109–130.

[17] W. Yang, X. Zhang, Y. Tian, W. Wang, J.-H. Xue, Q. Liao, Deep learning for single image super-resolution: A brief review, IEEE Transactions on Multimedia 21 (2019) 3106–3121.

[18] Y. Rivenson, Z. Göröcs, H. Günaydin, Y. Zhang, H. Wang, A. Ozcan, Deep learning microscopy, Optica 4 (2017) 1437–1443.

[19] H. Wang, Y. Rivenson, Y. Jin, Z. Wei, R. Gao, H. Günaydin, L. A. Bentolila, C. Kural, A. Ozcan, Deep learning enables cross-modality super-resolution in fluorescence microscopy, Nature methods 16 (2019) 103–110.

[20] T. Liu, K. De Haan, Y. Rivenson, Z. Wei, X. Zeng, Y. Zhang, A. Ozcan, Deep learning-based super-resolution in coherent imaging systems, Scientific reports 9 (2019) 1–13.

[21] K. de Haan, Z. S. Ballard, Y. Rivenson, Y. Wu, A. Ozcan, Resolution enhancement in scanning electron microscopy using deep learning, Scientific Reports 9 (2019) 1–7.

[22] J. Kennedy, R. Eberhart, Particle swarm optimization, in: Proceedings of ICNN’95-International Conference on Neural Networks, volume 4, IEEE, 1995, pp. 1942–1948.

[23] D. Arthur, S. Vassilvitskii, K-means++: The advantages of careful seeding, in: Proceedings of the Eighteenth Annual ACM-SIAM Symposium on Discrete Algorithms, SODA ‘07, Society for Industrial and Applied Mathematics, USA, 2007, p. 1027–1035.

[24] D. P. Kingma, J. Ba, Adam: A method for stochastic optimization, arXiv preprint 1412.6980 (2014).

[25] J.-Y. Zhu, T. Park, P. Isola, A. A. Efros, Unpaired image-to-image translation using cycle-consistent adversarial networks, in: Proceedings of the IEEE international conference on computer vision, 2017, pp. 2223–2232.

[26] G. Choi, D. Ryu, Y. Jo, Y. S. Kim, W. Park, H.-s. Min, Y. Park, Cycle-consistent deep learning approach to coherent noise reduction in optical diffraction tomography, Optics express 27 (2019) 4927–4943.

[27] M. Chen, N. McReynolds, E. C. Campbell, M. Mazilu, J. Barbosa, K. Dholakia, S. J. Powis, The use of wavelength modulated raman spectroscopy in label-free identification of t lymphocyte subsets, natural killer cells and dendritic cells, PLOS ONE 10 (2015) 1–14.

[28] N. McReynolds, F. G. M. Cooke, M. Chen, S. J. Powis, K. Dholakia, Multimodal discrimination of immune cells using a combination of raman spectroscopy and digital holographic microscopy, Scientific Reports 7 (2017).

[29] T. Peng, Detect circles with various radii in grayscale image via hough transform, Avaiable at https://www.mathworks.com/matlabcentral/fileexchange/9168-detect-circles-with-various-radii-in-grayscale-image-via-hough-transform (2020).

